# Analogue signaling of somato-dendritic synaptic activity to axon enhances GABA release in young cerebellar molecular layer interneurons

**DOI:** 10.1101/2022.10.18.512768

**Authors:** Federico F. Trigo, Shin-ya Kawaguchi

**Affiliations:** Departamento de Neurofisiología Celular y Molecular, Instituto de Investigaciones Biológicas Clemente Estable, Avenida Italia 3318, Montevideo 11600, Uruguay; Department of Biophysics, Graduate School of Science, Kyoto University Oiwake-cho, Kitashirakawa, Sakyo-ku, Kyoto, 606-8502, Japan

## Abstract

Axons are equipped with the digital signaling capacity by which they generate and faithfully propagate action potentials (APs), and also with the analogue signaling capacity by which subthreshold activity in dendrites and soma is transmitted down the axon. Despite intense work, the extent and physiological role for subthreshold synaptic activity reaching the axonal boutons has remained elusive because of the technical limitation to record from them. To address this issue, we made simultaneous patch-clamp recordings from the axonal varicosities of cerebellar GABAergic interneurons together with their parent soma or postsynaptic target cells in young rat slices and/or primary cultures. Our *tour-de-force* direct functional dissection indicates that the somatodendritic spontaneous EPSPs are transmitted down the axon for significant distances, depolarizing presynaptic boutons. These analogously transmitted EPSPs augment presynaptic Ca^++^ influx upon arrival of an immediately following AP through a mechanism that involves a voltage-dependent priming of the Ca^++^ channels, leading to an increase in GABA release, without any modification in the axonal AP waveform or residual Ca^++^. Our work highlights the role of the axon in synaptic integration.

## Introduction

Synaptic integration stands at the core of neuronal signaling. During synaptic integration, neuronal information provided by the presynaptic neurons is processed, leading to a new encoding of signaling that takes into account both the activity of the presynaptic neurons and the intrinsic properties of the integrating neuron. In the classical view of synaptic integration, the tasks of various compartments of the neurons are sharply defined: the somatodendritic compartment gathers information from presynaptic neurons; the axon initial segment sets the threshold for action potential (AP) firing; and the axon transmits the new AP to presynaptic terminals. In recent years, however, several studies have uncovered substantial deviations from this simple picture, and today it is clear that individual neurons do not necessarily behave as the « Platonic neuron » described by Coombs, Eccles and Fatt in the middle 1950s^1–3^.

This conceptual evolution was in part due to the description of a significant electrical coupling between somatic and axonal compartments in the subthreshold voltage range (termed “analogue signaling”). Although analogue signaling has been described in a variety of different preparations in mammals (for an exhaustive, recent review, see^4^), the quantification of the coupling with direct, simultaneous electrophysiological soma-axon recordings has been scarce in the literature because of the difficulties in recording from small varicosities of an intact axon in the majority of experimental preparations. As a corollary, analogue signaling has usually been studied by evaluation from indirect measurements and/or by strong subthreshold stimulation (using long, depolarizing or hyperpolarizing voltage changes), so that the incidence, extent and physiological role of analogue signaling for subthreshold spontaneous activity is only known in a handful of neuronal types^5–7^. Direct axonal patch-clamp recordings have shown, both in the hippocampal mossy fiber > CA3 synapse^5^ and in synapses between layer 5 pyramidal cells^6^, that subthreshold spontaneous or evoked somatodendritic activity can reach the axon. In cerebellar molecular layer interneurons (MLIs), on the other hand, previous data obtained by paired somatic recordings from pre- and postsynaptic neurons suggest that subthreshold coupling between the somatodendritic and axonal compartments is also substantial^8–12^, even for spontaneous activity^13,14^. However, the extent of such analogue signaling and its impact on synaptic outputs remain unclear because of the lack of a direct analysis by simultaneous recordings from pre- and postsynaptic structures in MLI synapses.

Cerebellar MLIs allow to quantitatively study the extent and functional impact of analogue signaling because their axonal bouton can be directly recorded with the patch-clamp technique, as shown by Southan and Robertson^15–17^ in pioneering, *tour-de-force* experiments. In the present work we performed for the first time simultaneous electrophysiological, whole-cell patch-clamp recordings from the soma and the axon of cerebellar MLIs in order to assess the coupling under regimes of physiological activity and understand its physiological role. We quantitatively show that in young MLIs (PN 13 to 17) the analog coupling of somato-dendritic synaptic activity to axon is substantial. By performing paired recordings from the axonal bouton and its postsynaptic target, we further show that this spontaneous synaptic activity coupled with an AP can affect transmitter release through a mechanism that involves the activation of voltage-dependent Ca^++^ channels, with no change in the presynaptic AP waveform or basal Ca^++^. Considered collectively with previous work, our findings highlight the importance of subthreshold coupling between neuronal compartments and further suggest that the process of synaptic integration is far more complex than classically envisaged, and that this is due to rich, largely unexplored signaling capabilities of the neuronal axon.

## Results

### Simultaneous soma-axon recordings in individual interneurons

In order to assess directly the degree of coupling between the somatodendritic and the axonal compartments we performed paired whole-cell recordings (WCR) from the soma and the intact axonal varicosities of cerebellar MLIs in an acute slice at near physiological temperature (34 °C). To do so we first patched the somatic compartment with an intracellular solution (IS) containing the fluorescent dye Alexa 594; after waiting for a few minutes for the dye diffusion we went on to patch an axonal varicosity with a pipette containing the same IS without Alexa dye. In the example shown in **Figure 1A** the distance between the center of the soma and the selected axonal varicosity is 150 μm (magenta line; minimum and maximum recording distances from the soma: 64.5 and 244 μm, respectively). **Figure 1B** shows a picture (the dotted rectangle in **A**) of the recording configuration during the cell-attached mode, a few seconds before rupturing the patch, and **Figure 1C** the same picture a few seconds after break-in (the intra-varicosity fluorescence disappears immediately after break-in). As shown in **Figure 1B** and **C**, the orthodromically transmitted axonal action potentials (axAPs) induced by a somatic depolarization were observed in cell-attached and whole-cell voltage and current clamp configurations: in cell-attached (**1B**); an unclamped spike current in voltage-clamp (VC; **1C, left**) and a short train of APs recorded in current-clamp (CC; **1C, right**).

**Fig. 1.**
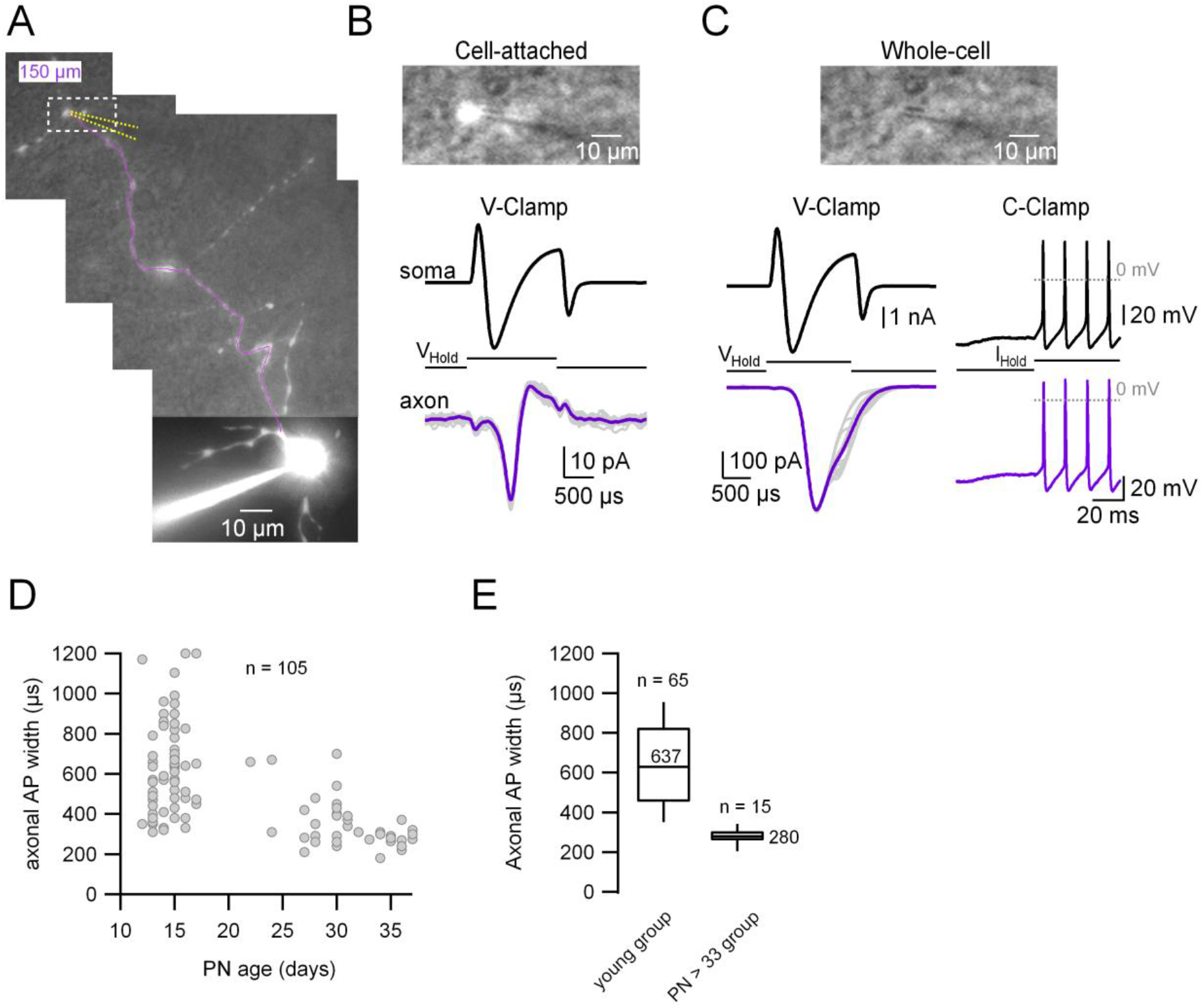
Simultaneous whole-cell recordings from the soma and its corresponding axonal presynaptic terminal. **A.** Fluorescence image of the recorded MLI. The patched varicosity is located at 150 μm from the soma. The position of the axonal patch pipette is represented with dotted yellow lines and the main axon in magenta. **B, C. Top.** Simultaneous transmitted and fluorescent light pictures of the area shown with the dotted rectangle in **A**, which shows the recorded axonal varicosity before (**B**) and after (**C**) rupturing the seal. In **B**, the fluorescence image is saturated and does not reflect the real size of the varicosity. **Bottom.** Somatic (upper) and axonal (lower), voltage- (left) or current-clamp recordings (right) of somatically induced Na^+^ current (or action potentials) and its propagation to the varicosity. **D.** AxAP width (measured from the cell-attached recordings) as a function of age (postnatal day 13 - 37). Total number of pairs = 105. **E.** Box plots of the 2 age groups in which the recordings were categorized. AxAP width of the immature age group (PN13 to 17): 639 ± 232 μs (median value 620 μs, n = 65). AxAP width of the mature age group (PN33 to 37): 278 ± 44 μs (median value 280 μs, n = 15). Here we set postnatal day 33 as the arbitrary limit between immature and mature animals.

Taking advantage of the cell-attached recording configuration presented above, which offers the possibility to characterize the axAP in unperturbed conditions^18^, we attempted to determine whether the AP width changes with development. As can be seen in **Figure 1D** and **E**, the width of the axAP decreases with age (PN13 to 17: 639 ± 232 μs; PN33 to 37: 278 ± 44 ms). We set postnatal day 33 as the arbitrary limit between immature and mature animals because the axAP width stabilizes from that age onwards. In addition, there was a clear decrease in the variability of the axAP width as well (CV of 0.3 for the younger age group; CV of 0.15 for the older group). Hereafter, all the experiments presented in the following sections were performed in the “young” age group as defined from the analysis of the axAP width.

### Substantial somato-axonal coupling for synaptic activity

To quantify the coupling for spontaneous synaptic activity we performed two types of experiments. In the first, we recorded spontaneous excitatory synaptic potentials (sEPSP) in paired somato-axonal recordings. **Figure 2A** shows the recording configuration (**top**) and transmitted and fluorescent light pictures of the recorded axonal varicosity (**bottom**; which in this case was contacting a Purkinje cell soma). As can be seen in **Figure 2B** and **C**, the simultaneous CC recordings from the soma and the axon show that every somatodendritic sEPSP is accompanied by the almost coincident appearance of an EPSP in the axonal varicosity (located here at 111.75 μm away from the soma). These axonal EPSPs are smaller in amplitude (mean ± SD amplitude of somatic and axonal EPSPs in this cell: 13.1 ± 2.0 mV and 7.8 ± 1.6 mV, respectively; n = 29 events, 50 s recording time, with an average coupling ratio [CR; see methods] of 0.6 ± 0.07), have a longer risetime (mean ± SD 10-90% risetimes of somatic and axonal EPSPs: 1.8 ± 0.49 ms and 5.7 ± 1.6 ms, respectively) and always appear later than the somatically recorded EPSPs (lag in the cross-correlation between the somatic and axonal recordings: 2.33 ms; **Figure 2D**). All of these data indicate that the somatically recorded EPSPs are closer to the source of current, namely the dendritic postsynaptic densities, and are compatible with classical cable models for the propagation of subthreshold events in neuronal compartments. The analysis of the decaying phase of the axonal and the somatic events shows that the axonal EPSP decay is well fitted with a mono-exponential function with a tau (20.4 ± 0.02 ms) which is extremely close to the slower time constant, τ2, of the somatic EPSP average (19.9 ± 0.02 ms). The fact that the final phase of decay, τ2, of the somatic and the axonal events is the same, is also in accordance with the prediction by cable models: the decay cannot be slower than the membrane time constant of the cell^19^.

**Fig. 2.**
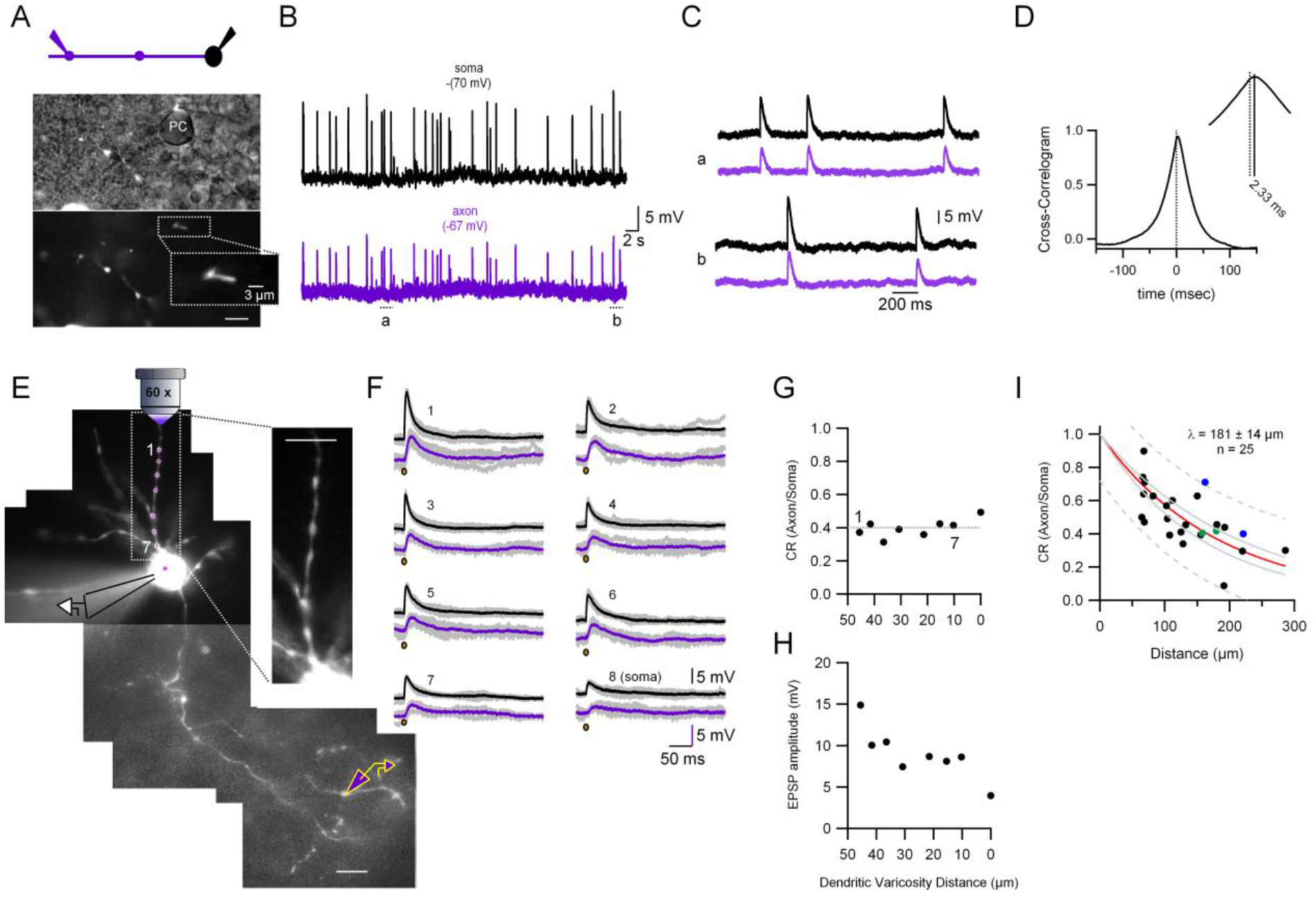
Quantification of the somato-axonal coupling for synaptic activity. **A. Top.** Recording configuration. **Bottom.** Pictures highlighting the position of the recorded axonal varicosity on top of a Purkinje cell somata during the cell-attached configuration. The inset shows an expanded view of the varicosity showing the fluorescence inside the patch pipette. **B.** CC recordings of spontaneous synaptic activity in the soma and the axon of the MLI. Resting membrane potential is indicated on top of each trace. **C.** Two selected time epochs are shown (**a** and **b** in **B**) with an expanded time-scale. **D.** Cross-correlogram of the traces shown in **B**. The inset shows the −20 to +20 ms interval. **E.** Fluorescent picture of an Alexa 594-filled MLI showing the experimental configuration. Glutamate was photolysed from MNI-glutamate at different dendritic varicosities (magenta spots 1 to 7 plus soma) while simultaneously recording from the soma and axon. The inset, which corresponds to a single picture, shows that all the dendritic varicosities tested are at the same depth, which ensures the same laser intensity. **F.** Somatic and axonal CC recordings of the depolarizations evoked by the photolysis of glutamate at the different dendritic varicosities (and soma) shown in **E**. Brown dots indicate the timing of the laser pulse. Laser pulse duration and power are 100 μs and 2 mW, respectively. **G.** CR between the soma and axon for each tested dendritic varicosity as a function of distance of the stimulated site to the soma. Dotted line is the average. **H.** Somatically recorded eEPSP amplitude as a function of distance. **I.** EPSPs amplitude ratios as a function of distance between somatic and axonal recording sites. The continuous red line shows the fit with an exponential function. Data include experiments with sEPSPs and eEPSPs. Individual points correspond to the average ratio calculated from the average of all the detected axonal and somatic EPSPs in each cell. Circles presented with the same color (blue and green) represent data from 2 different varicosities on the same axon. Dotted gray lines correspond to the prediction bands and solid green lines to the confidence bands.

In the second type of experiment, we performed spot laser photolysis of MNI-glutamate^20,21^. **Figure 2E** shows the recording configuration: MNI-glutamate was photolysed in 7 different dendritic varicosities plus the soma (**Figure 2E**; magenta spots). The laser-evoked EPSPs (eEPSPs) recorded simultaneously from the soma and the axon are shown in **Figure 2F**. As shown in **Figure 2G**, the soma-axon CR for eEPSPs at the different dendritic varicosities (and the soma) was not affected by the location of glutamate inputs at dendrites. The average CR (dotted line) was 0.4 ± 0.05 (mean ± SD). As a comparison, the CR for a DC voltage stimulus (resulting from 400 ms current injections in the soma) in this cell was 0.71 (not shown). **Figure 2H** shows the amplitude of the somatically recorded dendritic EPSPs as a function of distance. It can be seen that the higher amplitude is obtained when glutamate is photolysed at the farthest varicosity (and the lower amplitude when it is photolysed at the soma) which indicates that the number of postsynaptic glutamate receptors is probably larger at distant dendrites and that the amount of dendritic filtering in young MLIs is not prominent. In summary, glutamate photolysis in MLI dendrites to mimic sEPSPs confirms that dendritic synaptic potentials can travel long distances down the axon.

We next quantified the distance-dependence of the somato-axonal EPSP coupling by measuring the EPSP CR as a function of distance in 25 different soma-axon pairs (**Figure 2I**; both spontaneous and laser-evoked EPSPs were used for the analysis). The length constant, λ, of this distance-dependent relationship is 181 ± 14 μm (with a 95% confidence interval of 29 μm), which indicates that the analog coupling between the somatodendritic and axonal compartments in young MLIs is prominent for synaptic activity.

### Impact of analogue-digital coupling on the synaptic output and AP waveform in MLI boutons

It was shown before by our and other laboratories that analog signaling for long (> hundreds of ms) depolarizing pulses can affect release. We thus wondered whether the analogically traveling spontaneous synaptic activity lasting only tens of ms (see **Figure 2**) could affect release as well. In order to assess whether this was the case, we performed paired whole-cell recordings between MLIs and the postsynaptic Purkinje cells. Alexa 594 fluorescent dye was included in the IS to allow for MLI visualization (**Figure 3A**). Once a connection was established, the MLI was held in VC and two types of interleaved protocols applied: the control one consisted of a train of 5 stimuli (depolarization to 0 mV for 2 ms at 30 Hz) in order to induce somatic spikes; the test one was identical but the 1^st^ AP inducing pulse was preceded by a short (20 or 50 ms), subthreshold depolarization of the presynaptic MLI soma. From the PSC train (**Figure 3B**), we quantified both the PSC amplitude of the first response (**Figure 3C**) and the amplitude ratios of the subsequent PSCs in the train (**Figure 3D**). When the AP is preceded by a subthreshold depolarization the PSC amplitude is bigger than without the subthreshold depolarization (**Figure 3C**). Also, there is a decrease in the amplitude of subsequent PSCs (**Figure 3D**), which reflects an increase in the release probability at the 1^st^ stimulation and a lower availability of release-competent vesicles at the subsequent stimuli. These results indicate that short, somatically applied subthreshold depolarizations right before the AP can increase release, as revealed by an increase in PSC amplitude and the resultant decrease of later release.

**Fig. 3.**
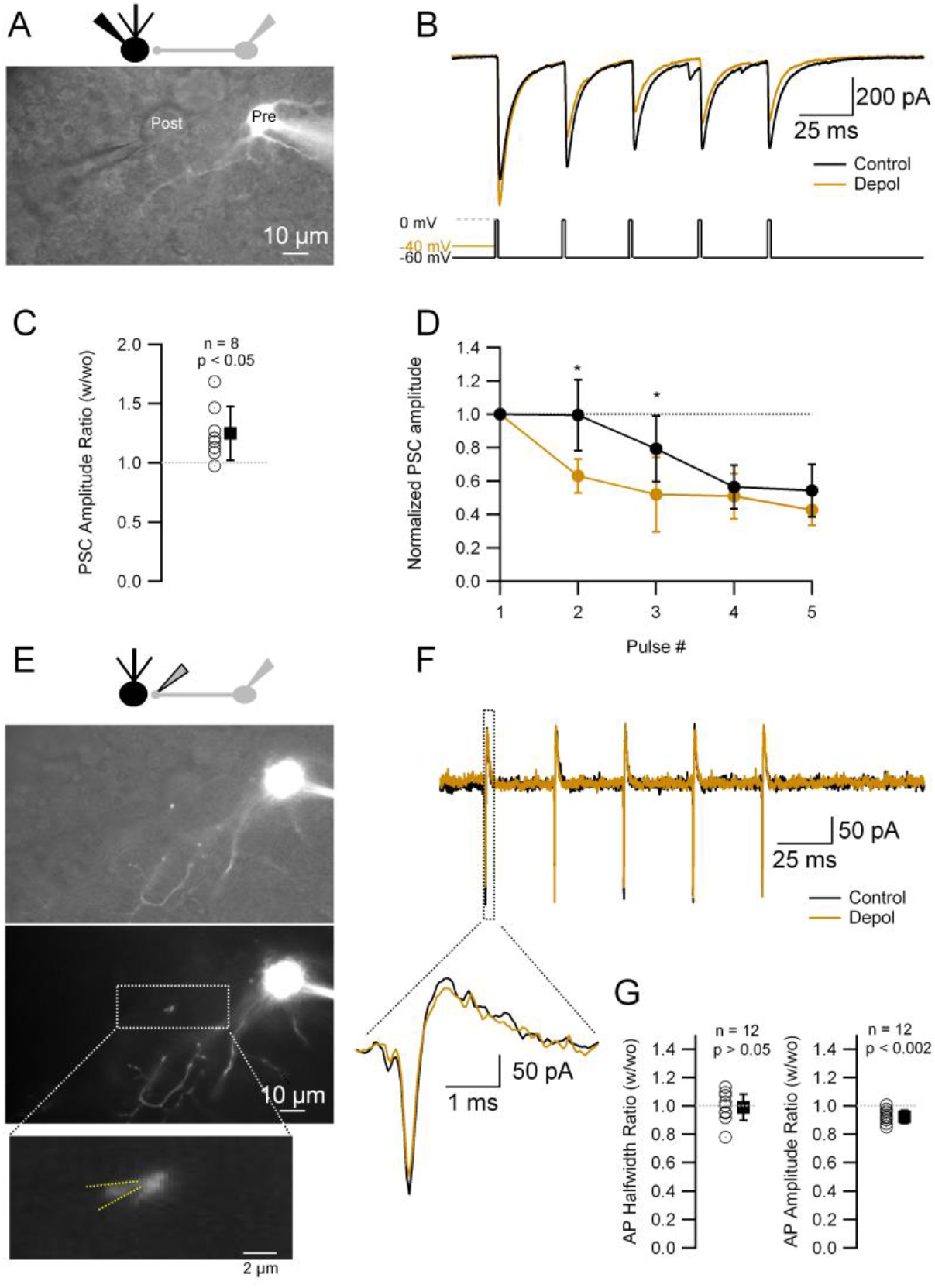
Short depolarizations increase PSC and short-term plasticity without any modification of the axAP width in cerebellar slices. **A. Top.** Recording configuration. **Bottom.** Superimposed transmitted and fluorescence light pictures of the synaptically-connected MLI and postsynaptic Purkinje cell. **B.** PSCs recorded from the Purkinje cell under two different experimental conditions: control (5 APs evoked in the MLI by depolarization to 0 mV from a HP of −60 mV at 33 Hz; black trace); test (same but with a 20 or 50 ms depolarization to −40 mV before the 1^st^ AP; brown trace). **C.** Ratio of the 1^st^ PSC in test over control conditions: 1.25 ± 0.23 (mean ± SD, n = 8, p < 0.05). **D.** Amplitudes of PSCs in a train normalized over the 1^st^ PSC in control and test conditions. Data show the averages ± SD from 7 cells. Asteriks indicate statistical significance (p < 0.05). **E. Top.** Recording configuration. **Middle.** Superimposed transmitted and fluorescence light pictures highlighting the MLI axonal varicosity on top of the Purkinje cell soma. Same pair as in A. **Bottom.** Fluorescence light picture of the MLI. The inset shows the details of the recorded axonal varicosity. Yellow dotted lines show the approximate position of the recording pipette. **F.** AxAPs recorded in the cell-attached configuration from the varicosity shown in **E** under the same experimental condition as in **B**. The inset shows an expanded view of the first axAPs in the train. **G.** Ratio of the 1^st^ AP widths (**Left**) and amplitudes (**Right**) in test over control conditions: mean width ± SD = 0.99 ± 0.09 (n = 12, p > 0.05); amplitude mean ± SD = 0.92 ± 0.06 (n = 12, p < 0.002).

Previous studies using longer depolarizations have proposed two primary mechanisms to explain the abovementioned phenomenon: a change in the axAP width or an increase in residual Ca^++^. By controlling the gating of presynaptic Ca^++^ channels, the duration of the axAP has a strong impact on the amount of released neurotransmitter^22,23^, and modulation of the axAP duration has been shown to happen as a result of analog transmission from the somatodendritic compartments^11,12^. In order to test whether these short, subthreshold depolarizations prior to an AP (preAP) modulate the axAPs we performed cell-attached recordings of the axonal varicosities. We first studied the axAP under the same experimental paradigm of somatic stimulation presented in **Figure 3A** to **D**. After finishing the Purkinje cell recording, we attempted to patch the presynaptic MLI axonal varicosity that was tentatively contacting the postsynaptic Purkinje cell. For that aim, the postsynaptic Purkinje cell pipette was removed and replaced by a smaller-tip electrode in order to perform a cell-attached recording of APs from the presynaptic cell axon. **Figure 3E** shows an example corresponding to the same MLI > Purkinje cell pair as in **Figure 3A**. After patching the MLI axonal bouton, the same (control and test) protocols were applied. Under these conditions, the axAP width remained unchanged between the control and test protocols (**Figure 3F** and **G**; 4 recordings with prior postsynaptic Purkinje cell recordings and 8 recordings where only soma-axon recordings were performed). The first axAP amplitude, on the other hand, showed a small, although significant, decrease (8%; **Figure 3F** and **G**), which is probably due to a decrease in the driving force for sodium influx during the AP onset because of the somatically-elicited depolarization arriving at the varicosity: given that the CR of DC signal is ~ 70 % as noted above, the Vm at an axonal varicosity would be 14 mV higher upon the 20 mV somatic depolarization, which is expected to decrease the driving force for sodium ions by about 10 %. Altogether, these experiments indicate that the change in synaptic efficacy elicited by short, EPSP-like subthreshold depolarization prior to an AP does not involve any change in the axAP width.

To further examine the stability of axAP in MLIs, we also explored the main features of the axAP under other paradigms of stimulation. One of the main conclusions of these experiments is that AP propagation from the soma to the axon is extremely reliable in MLIs; indeed, we never observed propagation failures even at the highest firing rates attained (500 Hz), independently of whether the recording was performed from the main axon or axonal collaterals. **Figure 4** shows a representative example of a paired “soma-bouton” recording where the soma is recorded in current-clamp and stimulated at increasing stimulation intensities, and the propagated AP recorded in the axon in the cell-attached configuration (**Figure 4A** and **B**). **Figure 4B** shows the instantaneous somatic firing frequency as a function of stimulus intensity for the 40 different trials. **Figure 4C** shows the raster plots for both the somatic and axonal spikes, which indicate that every somatic spike is accompanied by the corresponding axonal one, even at the higher firing frequencies (350 Hz in this example). **Figure 4D** and **E** show the half-width and amplitude, respectively, of the axAP plotted as a function of the cell’s firing frequency. It can be seen that the axAP half-width is extremely stable, even at the higher (> 300 Hz) firing frequencies (**4D**) while the axAP amplitude shows a small, although significant, reduction with frequency (**4E**). In order to further study the relationship between half-width and firing frequency, we designed a paired pulse experiment where the soma was stimulated twice at decreasing time intervals (**Figure 4F**). Even at the shortest intervals, the axAP width (2^nd^ over 1^st^ axAP) does not vary. **Figures 4G** and **H** shows the results from 11 different soma-axon pairs. The axAP amplitude was very stable as well, although a small decrease can be seen at the highest firing frequencies (**Figure 4H** and **inset**). This is in contrast to the somatically recorded AP, which shows a dramatic decrease in the amplitude at high but also low firing frequencies (black traces in **Figure 4A, F** and **H**). These results suggest that the AP is fully regenerated downstream of the AIS and they also show that the maximal firing frequencies that can be attained by stimulating the soma (**Figure 4A**) are much lower than the maximal firing frequencies that the axon can produce. In summary, these data indicate that the axAP in MLIs has a high safety factor in terms of the conduction reliability and is particularly impervious to activity-dependent modifications.

**Fig. 4.**
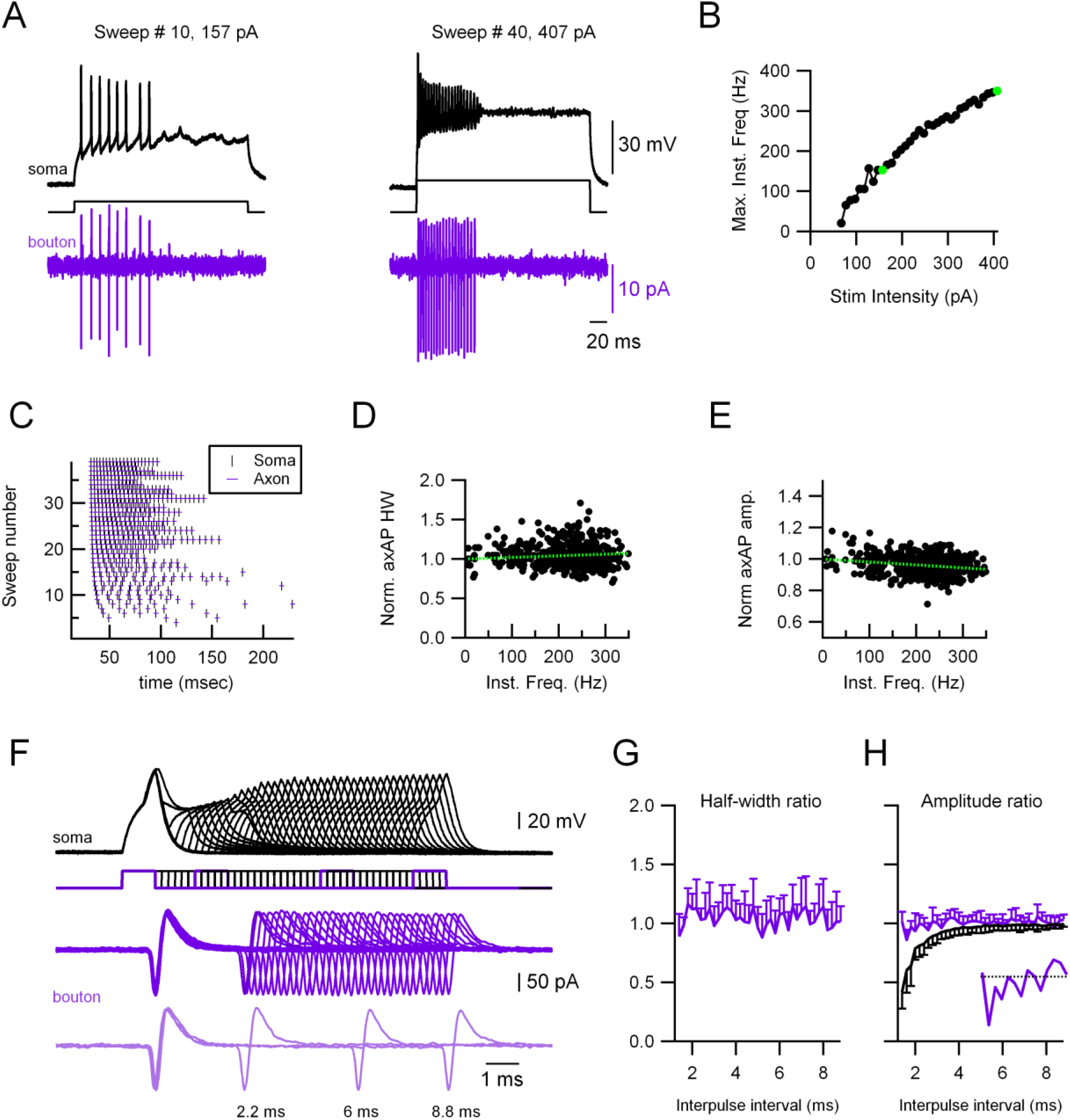
Highly reliable and stable axAP in MLI boutons. **A.** Representative experiment showing a paired recording between the soma (whole-cell current-clamp; black traces) and the axon (cell-attached in voltage-clamp; magenta traces). Firing was induced by injecting current through the somatic electrode (40 trials, 67 to 407 pA in 10 pA increments, 300 ms duration). **Left.** Responses at 157 pA stimulation intensity. **Right.** Responses at 407 pA stimulation intensity. **B.** Maximal firing frequency measured from the somatic recordings. The green dots correspond to the sweeps shown in **A**. **C.** Raster plots of the somatic and the axonal spikes for the 40 different trials, showing no axAP propagation failure. **D, E.** Normalized axAP width (**D**) and amplitude (**E**) as a function of instantaneous frequency. Green, dotted lines show a linear fit to the data. Spearman rank correlation test shows no significant correlation between axAP width and firing frequency (**D**) and a significant negative correlation between the axAP amplitude and firing frequency (**E**). The axAP widths and amplitudes are shown normalized to the first AP in each train to avoid errors due to fluctuations between trials. **F.** Representative experiment showing a paired recording between the soma (whole-cell current-clamp; black traces) and the axon (cell-attached in voltage-clamp; magenta traces). Firing was induced by injecting twin current pulses at varying intervals through the somatic electrode (each current pulse was 1 nA and 1 ms duration). Light magenta, lower traces, show a selection of 3 trials at different time intervals (8.8, 6 and 2 ms). **G, H.** Ratios (2^nd^ over 1^st^ axonal AP) of axAP half-width (**G**) and amplitude (**H**) as a function of interpulse interval. Magenta traces correspond to the axAPs and black trace to the somatically recorded APs (in current-clamp). Data corresponds to the averages of 11 experiments. Only either the positive or negative SD is shown for clarity. The inset in **H** shows the axAP amplitude ratio for the intervals 1.2 to 4 ms.

### EPSPs prior to an AP increase presynaptic Ca^++^ influx and GABA release

The results presented above (**Figure 3** and **4**) indicate that transmitter release from MLI terminals can be modulated by a subthreshold, EPSP-like potential changes in the absence of any change in the axAP width. To study how the synaptic outputs from MLI boutons are augmented without any change in the AP waveforms, we turned to the primary cerebellar culture preparation, where interneurons can be sparsely infected with an adeno-associated virus (AAV) that drives the expression of eGFP. This allows to do simultaneous recordings of both a single MLI axonal varicosity and its postsynaptic cell and hence to perform a detailed analysis of the release at the synapse.

In order to test whether small depolarizations similar to spontaneous EPSPs can affect release we performed paired recordings of a single varicosity and its postsynaptic partner. The presynaptic varicosity was voltage-clamped and stimulated by a voltage waveform that consisted of either a single AP (**Figure 5A**, black, left traces) or an AP preceded by two consecutive EPSPs (**Figure 5A**, brown, right traces; for details see methods). Upon the application of a realistic stimulus represented by the control AP waveform, a presynaptic calcium current (ICa^++^_pre_) is induced that triggers a PSC (**Figure 5A**). When the AP is preceded by the EPSPs the corresponding ICa^++^_pre_ is larger and so is the PSC. **Figure 5B** depicts the temporal relationship of the onset (dotted line “a”) and the peak (dotted line “b”) of the ICa^++^_pre_ in relation to the AP, which shows that the difference between the Ca^++^ currents in both conditions does not appear at the onset but at the peak of the current, during the decaying phase of the AP, suggesting that more Ca^++^ channels are activated by an AP coupled with preceding EPSPs. **Figure 5C** shows the relative increases (AP waveform with over without EPSPs) of the ICa^++^_pre_ and PSC amplitude, that are both significant. During the course of our experiments we also patched a cerebellar granule cell terminal contacting a MLI. The same experiments performed in such a pair did not show any increase in the preICa^++^ nor PSC amplitude (**supplementary Figure 1A**).

**Fig. 5.**
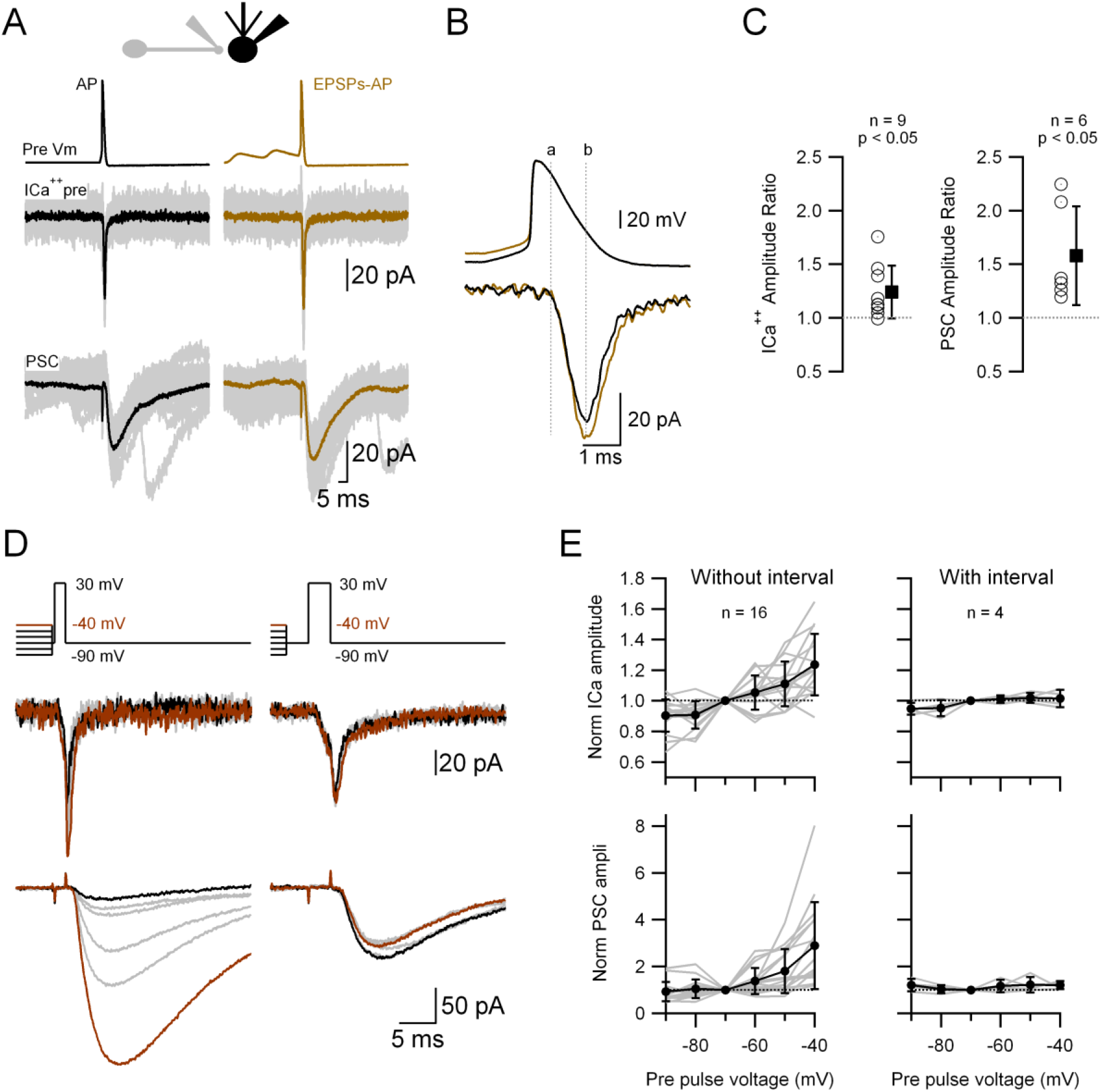
Passively propagated synaptic activity increases AP-induced ICa^++^ and PSC in cerebellar primary cultures. **A.** Simultaneous, VC recording of a presynaptic varicosity and the postsynaptic cell. Upper traces show the voltage waveforms applied, middle traces the recorded ICa^++^_pre_ and bottom traces the PSC. **B.** Expanded AP waveforms (upper traces) and the resulting ICa^++^_pre_ (bottom traces). The “a” and “b” dotted lines correspond to the onset and peak of the ICa^++^_pre_, respectively. **C.** Relative increases (waveform with EPSPs over that without EPSPs) of the ICa^++^_pre_ (**left**; 1.24 ± 0.25, n = 9, p < 0.05) and PSC (**right**; 1.58 ± 0.46, n = 6, p < 0.05) amplitudes. **D.** Simultaneous, VC recording of Ca^++^ influx into a presynaptic varicosity (middle traces) and the PSCs in its postsynaptic partner (lower traces), without (**left**) or with (**right**) a 3-ms interval between the small and large presynaptic voltage pulses. The upper traces show the voltage waveforms applied. The suprathreshold depolarizations lasted 1.5 ms (left) and 3 ms (right). **E.** Normalized ICa^++^_pre_ (upper graph) and PSC (lower graph) as a function of prepulse voltage for stimuli with (right) or without (left) a 3-ms interval between the prepulses and the suprathreshold depolarization.

To gain a better understanding of the relationship between the subthreshold potential and release, we performed a similar type of experiment to the one presented in **Figure 5D**, but now the ICa^++^_pre_ (and release) was triggered by a square depolarization (to 30 mV) preceded by 20 ms depolarizations to different voltages (−90 to −40 mV). **Figures 5D** and **E**, left, show that the ICa^++^_pre_ (and the subsequent PSC) is highly dependent on the subthreshold voltage before the suprathreshold pulse: the more depolarized is the presynaptic Vm, the larger are the ICa^++^_pre_ and subsequent release. A complete statistical analysis of the responses induced by the different prepulses is shown in **Supplementary Figure 2**, where the responses (ICa^++^_pre_ and PSC) obtained with the different prepulses were compared between each other. Again, there is no evidence of any activation of the ICa^++^_pre_ during the subthreshold depolarization. When a 3 ms interval is inserted between the subthreshold and suprathreshold pulses, the preceding subthreshold voltage levels do not have any influence on the ICa^++^_pre_ and transmission (**Figure 5D**, and **5E**, right). When the same protocol was applied to the granule cell terminal, there was no change in either of the 2 amplitudes (ICa^++^_pre_ or PSC; **Supplementary Figure 1B**). In summary, taken together these results strongly suggest that the subthreshold, short depolarization just before the AP impacts on the number of Ca^++^ channels that are activated, leading to dynamic changes of the efficacy of synaptic transmission.

### Impact of subthreshold depolarization on Ca^++^ channels in MLI boutons

To understand the mechanism by which the subthreshold depolarization increases the number of Ca^++^ channels activated upon the arrival of a subsequent AP, independently from any change in the axAP width, we recorded the ICa^++^_pre_ in a bouton and performed a biophysical characterization of it (kinetics and activation voltage) upon depolarizing pulses. When the terminal is voltage-clamped to 0 mV for various durations, an inward ICa^++^ develops with an activation time constant of around 3.0 ms and little or no inactivation (**Figure 6A**; n = 15). On the other hand, as shown in **Figure 6B**, the IV curve (from a total of 16 varicosities: 12 varicosities from primary cultures and 4 from slices) indicates that the take-off of the ICa^++^ is variable but always more depolarized than −40 mV, so it is unlikely that small depolarizations like the ones induced by EPSPs (as shown in **Figure 5**) would directly open the Ca^++^ channels, precluding the possibility of increased residual Ca^++^ by the subthreshold depolarization as the mechanism for the analog-digital coupling. Nevertheless, there still exists the possibility that a very small Ca^++^ influx may not be detected because buried into the recording noise. To test for this possibility, we analyzed the current variance (the variance is proportional to the number of open channels^24,25^) in presynaptic recordings at −70 mV and during depolarizations to −40 mV (the maximal depolarization tested; **Figure 6C**). This analysis shows that the variance during these 2 time epochs is the same, which is in agreement with the activation voltage shown on the IV curve before. These experiments confirm that the mechanism by which the preAP depolarization augments ICa^++^_pre_ and transmission does not involve any direct activation of Ca^++^ channels in the MLI bouton.

**Fig. 6.**
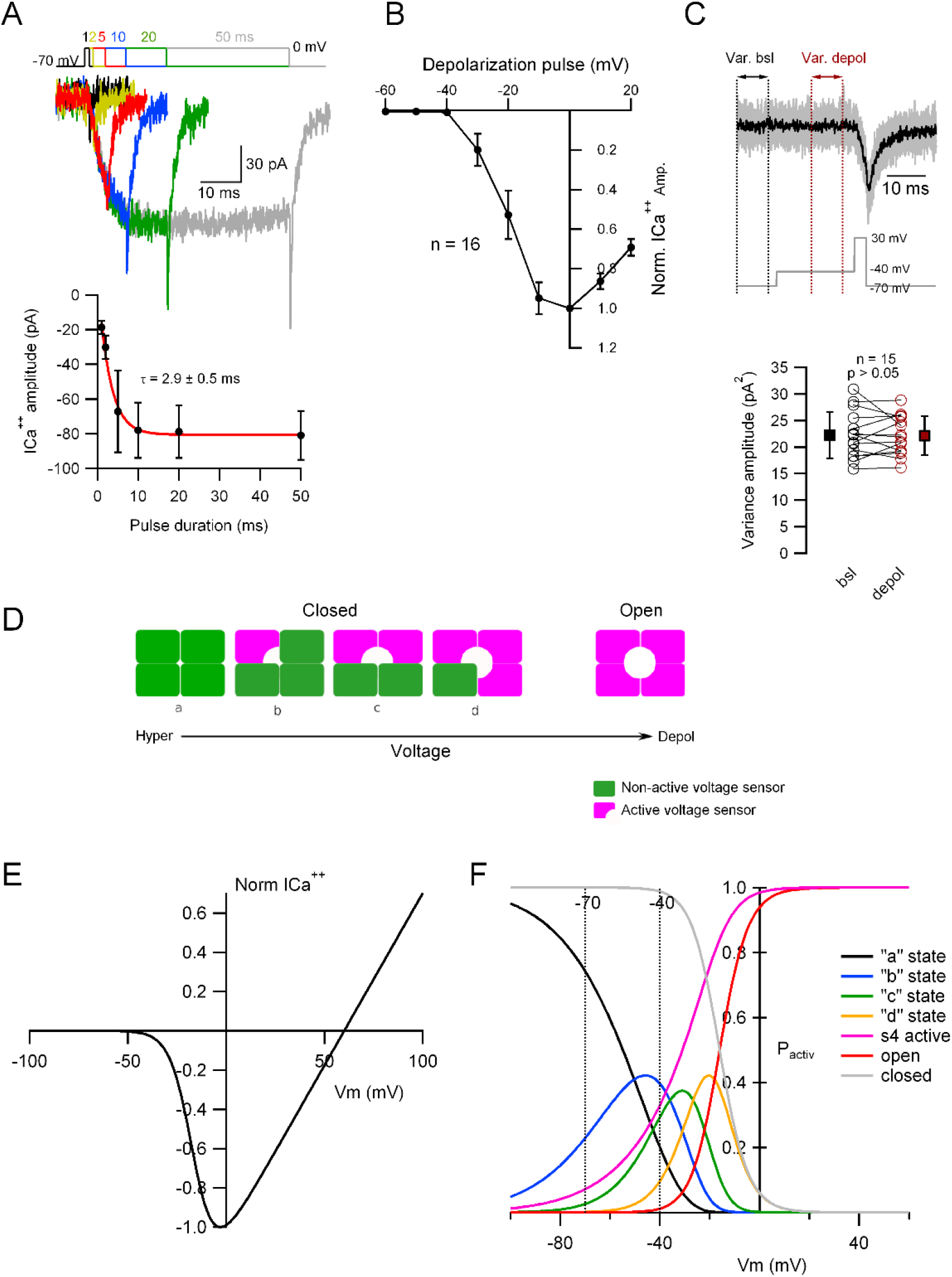
The preAP potential increases the probability of activating the voltage sensors of the Cav channels. **A. Top.** VC recordings showing representative traces of ICa^++^_pre_ upon depolarization to 0 mV for various durations (1, 2, 5, 10, 20 and 50 ms). **Bottom.** Average ICa^++^_pre_ as a function of stimulus duration (mean ± SEM). Red line is the fit to the data with an exponential function. Number of boutons analyzed: 1 ms = 4; 2 ms = 8; 5 ms = 12; 10 ms = 16; 20 ms = 16; 50 ms = 16. **B.** Current to voltage relationship of the ICa^++^_pre_ (mean ± SD; n = 16). **C. Top.** Representative ICa^++^_pre_ (upper traces) and the corresponding VC protocol (lower trace). The variance was measured in the time epochs indicated by the dotted lines. **Bottom.** Variance of individual cells (open symbols) and the mean ± SD (square, closed symbols) during baseline (−70 mV HP) and depolarization (−40 mV). Bsl variance is 22 ± 4.4 pA^2^ and during the depolarization 22 ± 3.6 pA^2^, n = 15 cells, p > 0.05. **D.** A simple model depicting the different configurations of the 4 voltage sensors of a voltage-gated Ca^++^ channel. When the membrane is depolarized (left to right; arrow) the probability of each of the 4 voltage sensors to go from the inactive to the active conformation increases. **E.** Simulated I/V relationship constructed from experimental data (**Figure 6B**). **F.** The probabilities to find the different states of the channel shown in **D** are plotted as a function of membrane potential (Vm). The dotted lines indicate the −70 and −40 mV Vm.

What is then the mechanism by which the analogue-digital coupling of EPSPs increases Ca^++^ influx and release in MLI boutons? Voltage-dependent Ca^++^ channels are composed of the pore-forming α1 subunit and other auxiliary subunits. The α1 subunit possess 4 repeats of structural assembly, each of them having 6 transmembrane segments including the voltage-sensing S4 domains and the pore-forming regions of the channel. The (non-conductive) closed state of the channel can assume 4 different conformations, “a” to “d”, represented in **Figure 6D**: the 4 voltage sensors are in their resting or non-activated position (a); there is 1 (b), 2 (c) or 3 (d) of the voltage sensors in their activated position. It is only when the 4 voltage sensors are simultaneously activated that the channel conducts (right scheme in **Figure 6D**). When we simply assume the Ca^++^ current activation characterized by the I-V curve as shown in **Figure 6E**, the probability of each voltage sensor being in its active position, P_s(v)_, varies with voltage and is calculated as the magenta trace in **Figure 6F**. The probability of the 4 voltage sensors being in their active position simultaneously (assuming that the individual probabilities are the same and that the 4 voltage sensors act independently; but see^26^), that is, the open conformation of the channel, can be represented as P_s(v)_^4^, and is characterized by the red trace in **Figure 6F** (which also shows the probabilities of states “a” to “d” and the total closed states of the channels [gray trace]). Considering the experiments presented in this study, the preAP depolarization (to −40 mV, vertical dotted line in **Figure 6F**) activates some of the voltage sensors, making the channel partly ‘primed’, so that upon the arrival of an AP the likelihood of having more channels shifting to fully open is higher than if departing from hyperpolarized potentials (for example −70 mV, vertical dotted line in **Figure 6F**). This simple mechanism relies exclusively on the operation of the voltage-gated Ca^++^ channels in the axonal bouton and is sufficient to explain the increase in GABA release presented here.

## Discussion

In this work we performed simultaneous patch-clamp recordings from the soma and axonal varicosities of cerebellar MLIs. These experiments allowed us to measure directly the synaptic activity coupling between the 2 compartments. As anticipated from previous works using other indirect methods (Ca^++^ imaging^9,10^, Ca^++^ and GABA photolysis^14^, voltage-sensitive dye imaging^11^ and electrophysiology^8,13^), we show here that the subthreshold voltage coupling in animals aged 12 to 17 days old is highly prevalent in these cells, even for short, subthreshold spontaneous activity (ie: EPSPs). We further show that the EPSPs that reach the axon can modulate presynaptic Ca^++^ influx and transmitter release through a mechanism that relies on the priming of the Ca^++^ channels by voltage, independently from any change in basal Ca^++^ or AP waveform.

### The mechanism by which orthodromic EPSP coupling affects release

It has been shown before by multiple groups, in cerebellar MLIs and in other neuronal types, that the somato-dendritic membrane potential can be transmitted down the axon and affects transmitter release through the modulation of voltage-dependent conductances^4^. This may be due to a change in the AP waveform (amplitude and/or duration^6,11^), a change of basal Ca^++^ concentration^9,10^ or to other, non-defined mechanisms^5,27^. The term analog-digital transmission is used to stress the fact that the subthreshold voltage (the analog signal) before the arrival of the AP (the digital signal) can affect release. The results presented here show that: a) the modulation of release does not require long or large voltage changes, but short and small, PSP-like voltage fluctuations of ≈ 10 mV can affect release; b) the mechanism involved does not include any change in the preAP waveform or in the presynaptic Ca^++^ influx right before the arrival of the AP. Even in these conditions, the subthreshold depolarization before the AP does induce an increase in the Ca^++^ influx during the AP and hence an increase in release. Our results are compatible with the following interpretation (schematized in **Figure 6**): during the subthreshold depolarization there is an increase in the voltage-dependent probability of the 4 voltage sensors of the Ca^++^ channels to go from the “resting-down” to the “activated-up” position^26^, but the pore of the channel remains closed. Once the AP is triggered, a bigger fraction of the voltage-gated Ca^++^ channels opens in relation to the control (no preAP depolarization) condition, where the probability of the voltage sensors to be in the activated position is at its minimum. The disappearance of the effect by a 3 to 5 ms interval after the subthreshold depolarization (**Figure 5D** and **E**) is compatible with this idea. In this sense, the mechanism described in this article constitutes, to the best of our knowledge, a new way by which somato-axonal analog coupling can modulate release. Although we cannot exclude that other mechanisms, as an increase in residual Ca^++^ or a change in the axAP width, may operate in other circumstances, they do not seem necessary to explain the effect described here. More importantly, these results highlight the relevance of recording directly from the axonal boutons in order to fully understand the mechanisms of synaptic release and AP propagation in CNS axons.

### Diverse analog-digital coupling mechanisms in MLIs

Subthreshold coupling between the somatic and axonal compartments in MLIs was described in the 90s by Glitsch and Marty^28^, who showed that long somatic depolarizations can increase GABA release through a pure analog transmission mechanism (similar to what happens in retinal amacrine cells). In 2011, Christie et al.^9^ and Bouhours et al.^10^ described for the first time an orthodromic subthreshold coupling in MLIs by which somatically-induced depolarizations are transmitted down the axon and increase AP-evoked release through an augmentation in basal Ca^++^, the so-called “analog-digital” transmission mechanism. It was later shown by Rowan and Christie^12^ that the analog-digital increase in GABA release is dependent on a modulation of the axAP width because of the inactivation of a K^+^ conductance (K_V_3). This is different from what was found by Bouhours et al., who showed that the analog-digital coupling was dependent on the activation of PKC. In a recent work, Blanchard et al.^29^ showed that the modulation of basal Ca^++^ with local Ca^++^ photolysis in a single MLI axonal varicosity can increase release through a modulation of the docking site occupancy. In the present work, both the amplitude and the half-width of the axAP are very resistant to various manipulations, like an increase in the stimulation frequency (**Figure 4**) and somatic depolarization (**Figure 3**), which is in agreement with what has been described by Alle and Geiger (see above) and by Ritzau-Jost et al.^18^, who showed that in neocortical (L5) fast-spiking GABAergic cells the width of the axAP does not change when the stimulation frequency increases. The fact that in our experiments the axAP width does not change, even when short prepulse depolarizations are applied before the AP (**Figure 3**), is in contrast to what has been found by Rowan and Christie^12^, who showed that somatic preAP depolarizations slow down the axAP. The difference may be due to a species difference (rat here vs mice in their work) or to the fact that the applied preAP depolarizations are longer in their study than those applied here.

In MLIs, it seems plausible that multiple mechanisms by which somato-axonal coupling affects release coexist, each one of them probably being initiated under a particular condition (for example, duration and amplitude of the somatic voltage changes). In this sense, it is interesting to note that the kinetics of the presynaptic ICa^++^ is slow (τ around 3 ms, **Figure 6A**). As a comparison, the ICa^++^ opens with an activation time constant of ~ 1 ms in Purkinje cell presynaptic boutons^30^ and ~ 2 ms in cerebellar granule cells^31^. This slow time constant of Ca^++^ channel activation in MLI boutons does not seem to be due to poor voltage clamp conditions. If this was the case, one would expect a correlation between the measured time constant and the Rs values: the higher the Rs, the slower the τ. However, no correlation was found between the τ and Rs values measured from the 16 different varicosities presented in **Figure 6A**. From the average Rs value calculated from those varicosities (83 MΩ), on the other hand, an approximate voltage clamp time constant of 200 μs can be calculated, assuming a varicosity capacitance of 1 to 2 pF. In summary, the slow τ of the ICa^++^ indicates that a very small fraction of the available Ca^++^ channels opens upon the arrival of an AP in MLIs, leaving a big verge for different modulation mechanisms to occur. Indeed, unlike MLI boutons, a cerebellar granule cell bouton does not show an augmentation of ICa^++^_pre_ and transmission upon pre-pulse depolarization before AP (**Supplementary Figure 1**).

### Ideas and speculation: The physiological role of somatodendritic spontaneous activity somato-axonal coupling

Our preliminary, non-published results suggest that the somato-axonal coupling efficiency for spontaneous activity that we describe in this study decreases with age. This implies that the increased GABA release may have some role in the development of synaptic contacts. Indeed, our experiments have been performed at postnatal days 13 to 17, a time-period where GABAergic synapses between MLIs and their postsynaptic partners (Purkinje cells and other MLIs) are still under maturation^32^. Interestingly, this time-period coincides with the time period where axonal GABA_A_ auto-receptors exert an facilitating effect on GABA release at the MLI boutons^33^. In the light of this observation, it is tempting to speculate that both the somato-axonal coupling of spontaneous activity and the GABA_A_ auto-receptors are part of the same activity-dependent mechanisms that contribute to the maturation of GABAergic contacts^34^. In more general terms, analogue-digital coupling may be the common physiological behavior of short-axon neurons, at least at early stages of neuronal development.

## Supplementary figures

**Figure S1.**
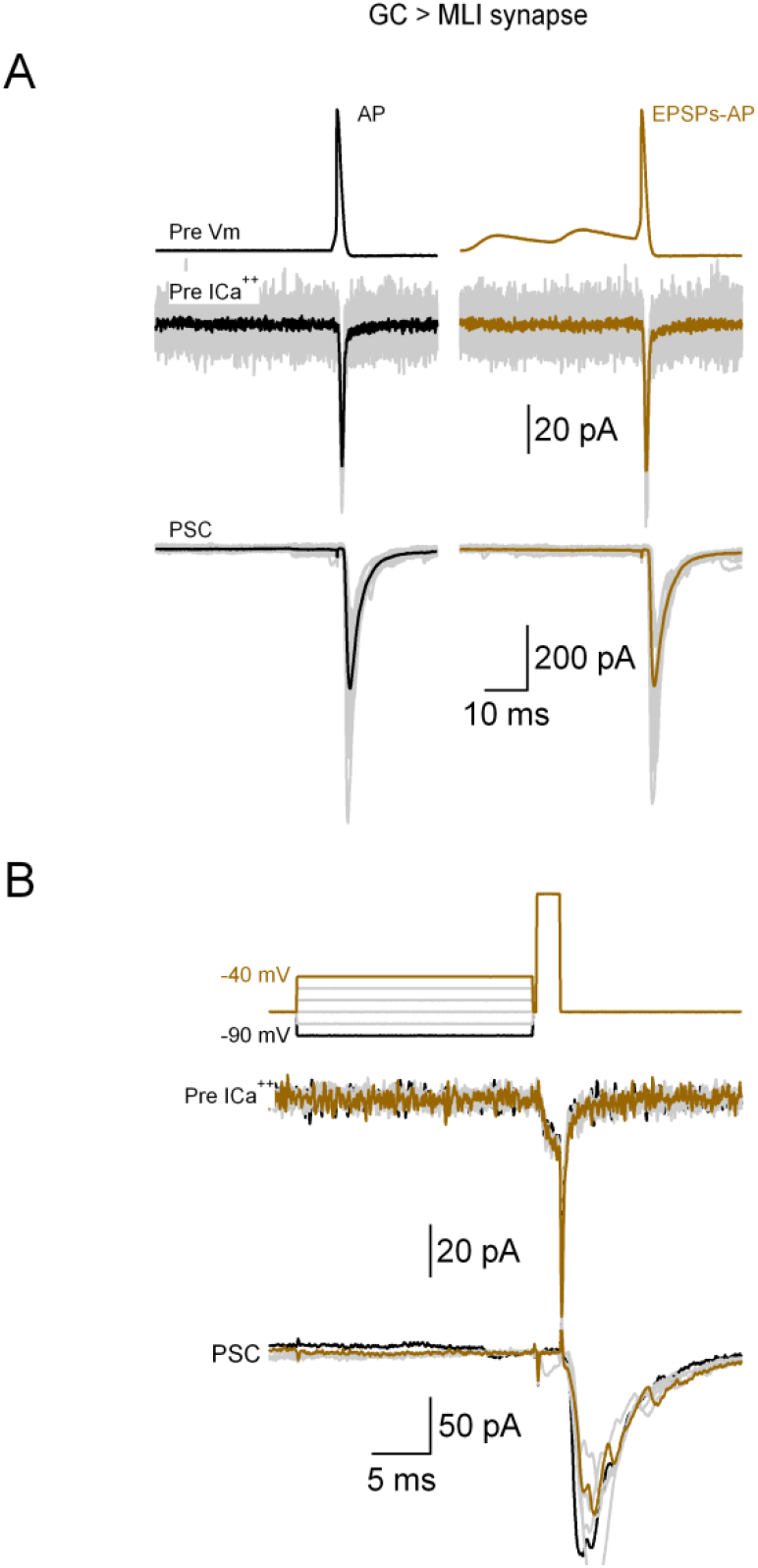
**A.** Simultaneous, VC recording of a presynaptic granule cell varicosity and its postsynaptic MLI. Upper traces show the voltage waveforms applied, middle traces the recorded preICa^++^ and bottom traces the PSC. **B.** Simultaneous, VC recording of the same granule cell. Middle traces show the preICa^++^ and the bottom traces the PSCs in the postsynaptic MLI. The upper traces show the voltage waveforms applied.

**Figure S2.**
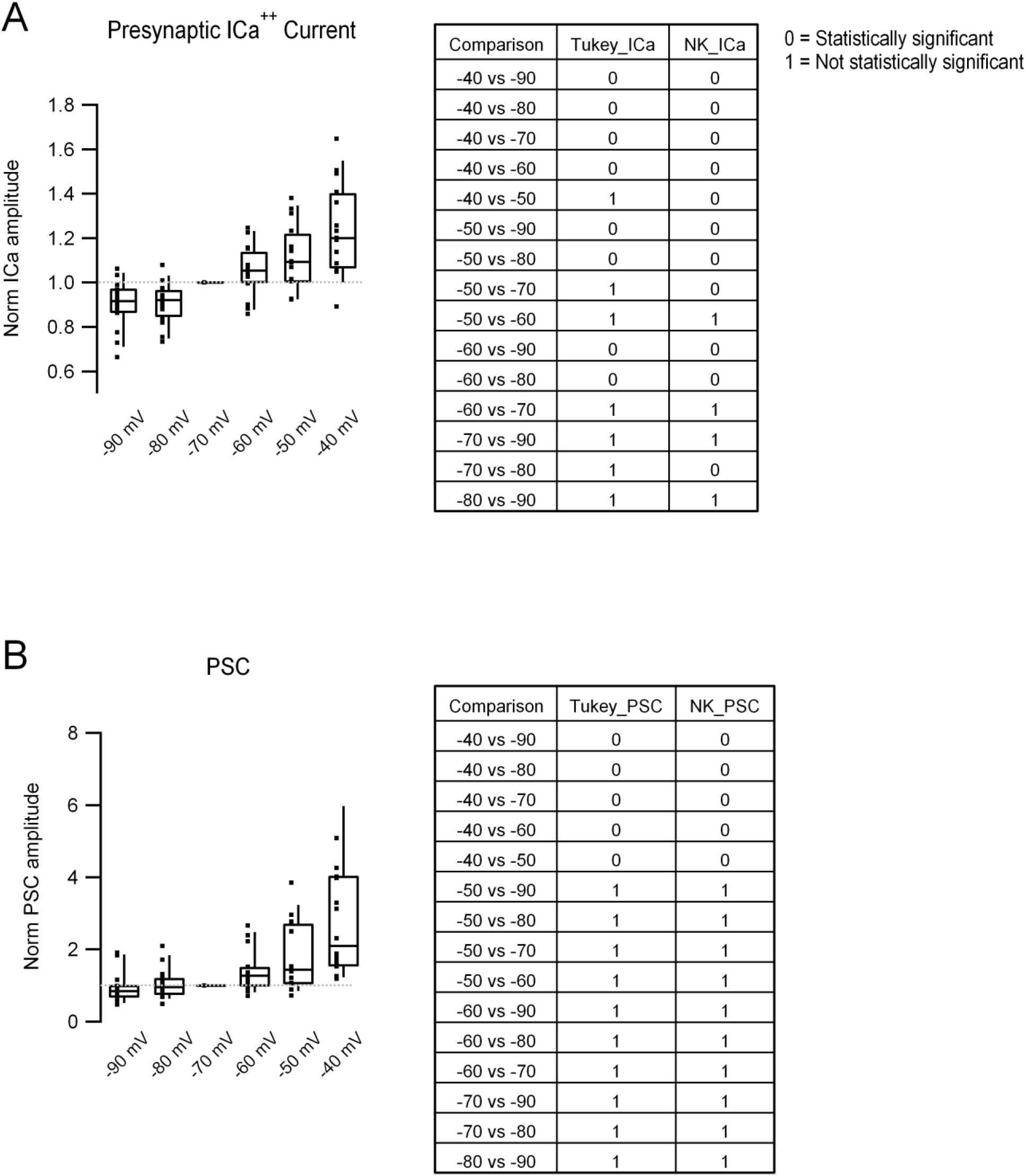
**A. Left.** Box plots showing the preICa^++^ amplitude obtained with the different prepulses. In each column, middle line corresponds to the median, upper and lower horizontal lines to percentiles 75 and 25 respectively, and upper and lower extremes to percentiles 90 and 10, respectively. Filled squares correspond to individual measurements. **Right.** Table showing the statistical comparison (Tukey, column 2, and Newman-Keuls, column 3, tests) of the different values. 0 corresponds to a statistically significant difference (rejection of the null hypothesis), and 1 to no difference (acceptance of the null hypothesis). **B. Left** and **Right.** Same analysis as in **A** but for PSC amplitude. Data corresponds to the 16 different pairs shown in F**igure 5D**.

## Materials and Methods

### Preparation of cerebellar slices

Slices were prepared from Sprague-Dawley rats aged 12 to 37 days old. After decapitation, the cerebellum was quickly removed in an ice-cold extracellular solution (ES), the cerebellum taken out and sagittal cerebellar slices (202 μm width) cut with a Leica vibroslicer (VT1200S). The slices were kept in a recovering chamber at 34 °C until use. The composition of the ES was, in mM: NaCl 115, KCl 2.5, NaH_2_PO_4_ 1.3, NaHCO_3_ 26, Glucose 25, Na-pyruvate 5, CaCl_2_ 2, MgCl_2_ 1, pH 7.4 when bubbled with carbogen (95 % O_2_ and 5 % CO_2_). For animals aged 12 to 21 days the same afore-mentioned ES was used for the recordings. For animals aged above 21 days old a KGluconate-based ES was used for the dissection^35^; its composition (in mM): KGluconate 130, KCl 15, EGTA 0.05, Hepes 20, Glucose 25 and DAP5 50 μM, pH 7.4 and bubbled with carbogen.

### Preparation of cerebellar cultures

The method for preparing primary dissociated cultures of cerebellar neurons was similar to that in a previous study^36^. Inhibitory interneurons were transfected with eGFP at 1 day after culture with adeno-associated virus (AAV) vector serotype 9 under the control of the CMV promoter. Interneurons could be visually identified from eGFP fluorescence. An axon of interneuron surrounding a PC soma was selected for whole-bouton recordings. Experiments were performed 3-5 weeks after preparation of the culture.

### Electrophysiology

Both the soma and axon of MLIs were recorded with the patch-clamp technique^37^ either in voltage or in current-clamp with a HEKA amplifier (EPC-10, double) and either Luigs & Neumann or Sutter manipulators. For the experiments presented in **Figures 1** to **4** a KGluconate-based IS of the following composition (in mM) was used: 165 KGluconate, 10 HEPES, 1 EGTA, 0.1 CaCl_2_, 4.6 MgCl_2_, 4 Na2ATP, 0.4 NaGTP, pH 7.3 and osmolarity 300 mOsm/KgH_2_O. Alexa 594 (0.04 mM) was also added to the somatic recording pipette. With this IS the somatic pipettes had resistances of around 6 MΩ and the axonal ones 25 MΩ. To record the Ca^++^ currents (**Figures 5** and **6** and **Supplementary figures**) a CsCl-based IS of the following composition (in mM) was used: 152 CsCl, 0.5 EGTA, 10 HEPES, 10.5 CsOH, 2 ATP, 0.2 GTP, pH 7.3 and osmolarity 300 mOsm/KgH_2_O. With this IS the somatic pipettes had resistances of around 4 MΩ and the axonal ones 18 MΩ. In these experiments TEA (2 mM) and TTX (200 nM) were added to the extracellular solution in order to block K^+^ and Na^+^ voltage-dependent conductances.

Somatic and axonal pipettes were pulled with either HEKA (Pip 6) or Narishige (PP-83) vertical pullers. For experiments in the slice, we first recorded from the soma with the Alexa 594 containing IS. After a 5 to 10 minutes waiting time the fluorescence illumination was turned on in order to identify the axon and a suitable, superficial varicosity. In order to patch the axon a second pipette containing the KGluconate-based IS (without Alexa) was used. The axon was patched by looking at the image created by the camera with both the bright field and fluorescent lights on (**Figures 1A** and **1B**). Pictures were taken at 1 Hz frequency in order to avoid photodamage.

Recordings in slices were done in an upright Olympus BX51W equipped with a 60×, 1.0 numerical aperture objective (NA). Experiments were done at near-physiological temperature (≈ 34 °C) with a Peltier system (Luigs & Neumann). Electrophysiological recordings from primary cultures were done in an inverted Olympus IX71 microscope equipped with a 40× objective (NA. 0.95). For experiments in slices, epifluorescence excitation was by light-emitting diode (LED) controlled by an OptoLED system (Cairn Research) at 572-/35-nm excitation and at 630/60 nm emission. Filters and dichroic were from Chroma Corporation (Vermont, USA). For experiments in cultures epifluorescence was by an LED light (Light Engine SOLA, Lumencore) at 450-/40nm excitation and 510-/50 nm emission; Fluorescent images were taken with Andor cameras (EMCCD Andor Ixon or a sCMOS Zyla 4.2).

Salts were either from Sigma Aldrich or Nacalai Tesque (Japan).

### Photolysis

Glutamate was photolysed from MNI-Glutamate (4-Methoxy-7-nitroindolinyl-caged-L-glutamate; Tocris Biotechne, UK) with a 405 nm laser (Obis, Coherent, USA) following well-established procedures^21^. Briefly, MNI-Glutamate was added directly to the bath at a final concentration of 500 μM and photolysed with 100 to 200 μs and 1 to 3 mW laser pulses. Duration and power of the laser pulses were adjusted to obtain laser evoked EPSPs similar to spontaneous ones in terms of amplitude and time course. It is well known that laser power decreases exponentially with depth^21^. In order to be certain that the same amount of glutamate was released at each tested dendritic varicosity, only dendritic uncaging sites localized at the same imaging plane were chosen (see inset in **Figure 2E**).

#### AP waveform

The voltage waveforms used in the experiments presented in **Figure 5A** (either the AP alone or the AP preceded by two EPSPs) were constructed from real AP and EPSPs recorded from an interneuron axonal varicosity. The width of the AP waveform used, 780 μs, was compatible with the width recorded in the cell-attached configuration (**Figure 1 D** and **E**). The resting Vm of the AP waveform is −70 mV and the peak of the AP is at 40 mV; the EPSPs interval is 19.5 ms, the EPSPs amplitude −58 (#1^st^ EPSP) and −53 mV (2^nd^ EPSP) and the duration of the depolarization (time between the onset of the EPSP and the onset of the AP) is 40 ms.

### Analysis

EPSPs were detected using TaroTools extensions in IgorPro (https://sites.google.com/site/tarotoolsregister/), and the selection of each event was visually confirmed before subsequent analysis. The EPSPs coupling ratio, CR, represents the ratio of the peak axonal and peak somatic depolarizations.

The distance between the center of the soma and the recorded varicosity was measured offline from reconstructions of the recorded cell with Fiji^38^. The axAP widths and amplitudes (**Figures 1, 3** and **4**) in cell-attached recordings were measured from peak to peak, which correspond to the maximal slopes of the rising and decaying phases of the AP voltage waveform.

### Statistics

Data are presented as mean ± SD unless otherwise stated. Statistical significance was tested with the Wilcoxon signed-rank test for paired and the Wilcoxon-Mann-Whitney test for non-paired data. The difference between groups was considered significant when p < 0.05. In **Figure 4** (axonal AP width and amplitude vs firing frequency), the correlation was assessed by the Spearman rank correlation test. In **supplementary Figure 2**, statistical significance between the means of the different groups was assessed with the Tukey and Neuman Keuls post-hoc tests.

## Acknowledgments

The authors thank Alain Marty for the critical reading of the manuscript.

## Funding

Agence Nationale pour la Recherche (ANR) JCJC grant ANR-17-CE16-0011-01 (FFT).

International Brain Research Organization (IBRO) Return Home Fellowship (FFT).

Japan Society for Promotion of Science, Core-to-Core Program A (SK).

Japan Society for Promotion of Science, KAKENHI grants 22H02721 and 22K19360 (SK).

Takeda Science Foundation (SK).

Naito Foundation (SK).

## Author contributions

FFT and SK participated in each step of the work and contributed equally to the paper. Correspondence should be addressed to either of the two authors.

## Data and materials availability

The data that support the findings of this study are available from the corresponding authors upon request.

